# Increased excitation-inhibition ratio stabilizes synapse and circuit excitability in four autism mouse models

**DOI:** 10.1101/317693

**Authors:** Michelle W. Antoine, Philipp Schnepel, Tomer Langberg, Daniel E. Feldman

## Abstract

Distinct genetic forms of autism are hypothesized to share a common increase in excitation-inhibition (E-I) ratio in cerebral cortex, causing hyperexcitability and excess spiking. We provide the first systematic test of this hypothesis across 4 mouse models (*Fmr1^−/y^*, *Cntnap2^−/-^*, *16p11.2^del/+^*, *Tsc2^+/-^*), focusing on somatosensory cortex. All autism mutants showed reduced feedforward inhibition in layer 2/3 coupled with more modest, variable reductions in feedforward excitation, driving a common increase in E-I conductance ratio. Despite this, feedforward spiking, synaptic depolarization and spontaneous spiking were essentially normal. Modeling revealed that E and I conductance changes in each mutant were quantitatively matched to yield stable, not increased, synaptic depolarization for cells near spike threshold. Correspondingly, whisker-evoked spiking was not increased *in vivo*, despite detectably reduced inhibition. Thus, elevated E-I ratio is a common circuit phenotype, but appears to reflect homeostatic stabilization of synaptic drive, rather than driving network hyperexcitability in autism.

## Introduction

Autism spectrum disorders (ASD) are a family of neurodevelopmental disorders characterized by social and communication deficits, restricted and repetitive behaviors or interests, and abnormal sensory responses (Geschwind 2009). ASD is highly genetically heterogeneous, with >100 identified risk genes that have diverse functions in transcriptional regulation, protein synthesis and degradation, synapse function and synaptic plasticity. However, whether genetically distinct forms of ASD share a common dysfunction at the neural circuit level remains unclear.

One long-standing model is that genetically distinct forms of ASD share a common increase in synaptic excitation to inhibition (E-I) ratio in cerebral cortex, which drives hyperexcitability, excess spiking and increased noise in cortical circuits. This is hypothesized to cause the cognitive and behavioral symptoms of autism (Nelson and Valakh, 2015, Rubenstein and Merzenich, 2003). Prior synaptic physiology studies using transgenic mouse models of ASD provide mixed support for this E-I ratio hypothesis. Many report reduced inhibition (Chao et al., 2010; Gibson et al., 2008; Han et al., 2012; Liang et al., 2015; Mao et al., 2015; Wallace et al., 2012), often coupled with a smaller decrease in synaptic excitation (Gibson et al., 2008, Mao et al., 2015,Wallace et al., 2012). However, others report a greater decrease in excitation than inhibition (Dani et al., 2005, Delattre et al., 2013, Unichenko et al., 2017,Wood and Shepherd, 2010), or increased inhibition (Dani et al., 2005, Harrington et al., 2016,Tabuchi et al., 2007). Variation across studies in brain area, cell type, ASD genotype, and physiological methods prevent identification of common synaptic and local circuit defects in ASD.

Critically, whether increased E-I ratio yields hyperexcitable cortical networks in ASD remains fundamentally unclear. From basic biophysics, increased E-I conductance ratio does not necessarily lead to stronger synaptic depolarization or spike probability. Empirically, some ASD mouse models show increased cortical spiking activity in vivo (Peixoto et al., 2016, Rotschafer and Razak, 2013,Zhang et al., 2014), but most show no or modest abnormalities (Dolen et al., 2007, Goncalves et al., 2013, O’Donnell et al., 2017, Wallace et al., 2017) or even reduced cortical activity (Banerjee et al., 2016, Durand et al., 2012, Garcia-Junco-Clemente et al., 2013,Unichenko et al., 2017). In humans, increased network excitability is suggested by increased seizure prevalence in some forms of ASD, but seizures only occur in a subset of patients and EEG may be normal between seizures when ASD symptoms present (Samra et al., 2017,Tuchman et al., 2010). Many ASD mouse mutants show clear behavioral phenotypes in the absence of elevated cortical activity, spontaneous seizures or abnormal EEG (Dhamne et al., 2017, Goorden et al., 2007, Peñagarikano et al., 2011,Thomas et al., 2017). Thus, whether E-I ratio is systematically altered across genetically distinct forms of autism, and whether this drives excess spiking in cortical circuits, remain unclear. Optogenetic manipulations of E-I ratio and spiking in prefrontal cortex induce and ameliorate social behavioral deficits, but this doesn’t mean that elevated E-I ratio or excess spiking is the endogenous cause of social impairment in ASD mice (Yizhar et al., 2011,Selimbeyoglu et al., 2017).

We tested for common circuit defects in somatosensory cortex (S1) of four genetically distinct, well-validated mouse models of ASD (*Fmr1^−/y^*, *Cntnap2^−/-^*, *16p11.2^del/+^*, *Tsc2^+/-^*). S1 is a reasonable focus because tactile disturbances are common in ASD (Robertson and Baron-Cohen, 2017), and S1 excitatory and inhibitory circuits are well characterized. We studied the feedforward circuit from layer (L) 4 to L2/3 pyramidal (PYR) cells, which is the first step in intracortical sensory processing. L4-L2/3 feedforward excitation and inhibition are integrated by PYR cells to evoke sparse spiking. L4-L2/3 feedforward inhibition is mediated by parvalbumin (PV) interneurons, which are implicated in ASD (Selten et al., 2018). We systematically tested each ASD mutant *in vitro* and *in vivo* for abnormal synaptic excitation and inhibition in L2/3 PYR cells, abnormal network spiking, and impaired sensory coding. *Fmr1^−/y^* mice have impaired inhibition in L4 (Gibson et al., 2008), and *Cntnap2^−/-^* mice have fewer PV interneurons (Vogt et al., 2017), but E-I ratio phenotypes in L2/3 are unknown in any of these mutants. Thus, these four mutants provide a strong test for general applicability of the E-I ratio hypothesis.

We found that all ASD mutants exhibited decreased inhibition and more weakly decreased excitation, yielding increased E-I conductance ratio. However, contrary to the E-I ratio hypothesis, synaptic conductance modeling showed that these E-I changes were quantitatively matched to preserve peak synaptic depolarization, not increase it. Consistent with this, peak synaptic depolarization and spiking were remarkably normal in ASD mutants, *in vivo* and *in vitro*. Thus, rather than promoting circuit hyperexcitability, increased E-I ratio appears to be a compensatory mechanism that stabilizes synaptic depolarization and spiking excitability in these ASD genotypes.

## Results

### L4-L2/3 synaptic currents and E-I conductance ratio

We tested for abnormal synaptic currents in S1 slices from juvenile *Fmr1^−/y^*, *Cntnap2^−/-^*, *16p11.2^del/+^* and *Tsc2^+/-^* mice and age-matched wild type controls. We first measured L4-evoked feedforward excitatory and inhibitory currents (EPSCs and IPSCs) converging onto single L2/3 PYR cells (**Figure 1A**). EPSCs and IPSCs were separated in whole-cell voltage clamp by holding at −72 and 0 mV, the reversal potentials for excitation and GABA-A inhibition. L4-evoked IPSCs were blocked by NBQX and D-APV (to 2.7±1.6% of control, n=3 cells), and thus represent disynaptic feedforward inhibition. For each PYR cell, we found the minimum L4 stimulation intensity required to evoke a detectable EPSC, denoted Eθ, and then measured input-output curves for EPSCs and IPSCs at 1.0-1.5x Eθ. For analysis, currents were integrated over 20 ms, matching the time scale of L2/3 sensory integration *in vivo* (McGuire et al., 2014). Stimulation at Eθ generally evoked small EPSCs and IPSCs. Increasing stimulus intensity recruited steadily larger EPSCs and disproportionately larger IPSCs, so that inhibition dominated at ≥ 1.2x Eθ, as in prior studies (House et al., 2011,Xue et al., 2014). Example cells for all genotypes are shown in **Figure 1B**.

**Figure 1:**
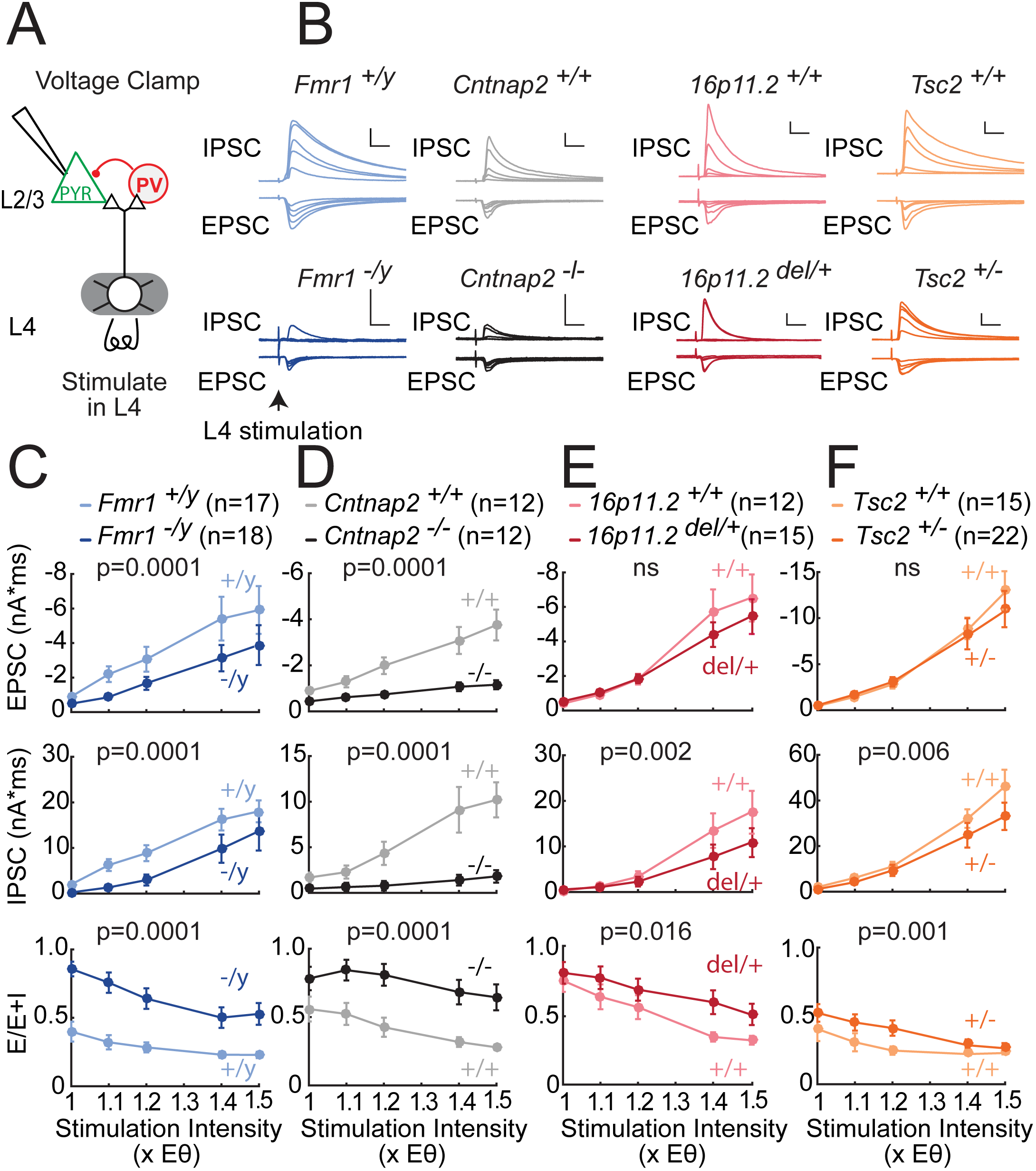
Deficits in feedforward excitatory and inhibitory synaptic conductances in L2/3 PYR cells in four ASD mouse lines. **(A)** Configuration for measuring L4-L2/3 feedforward EPSCs and IPSCs in L2/3 PYR neurons in S1 slices. **(B)** L4-evoked EPSCs and IPSCs at 1.0, 1.1, 1.2, 1.4 and 1.5x Eθ from 8 example L2/3 PYR cells from ASD mutant mice (darker colors) and corresponding wild types (lighter colors). Scale bars: 10 ms, 500 pA. **(C-F)** Average input-output curves for EPSCs, IPSCs, and E-I conductance ratio calculated as E/(E+I). Plots show mean ± SEM across cells. P-values are for genotype factor in a 2-way ANOVA on log-transformed data. N, number of cells.

*Fmr1^−/y^* mutants had smaller EPSCs than *Fmr1^+/y^* wild types (**Figure 1C**; n=17, 18 cells, p=0.0001, two-factor ANOVA on log-transformed data). Mouse N’s for all slice physiology measurements are in **Table S1**. IPSCs were also reduced strongly in *Fmr1^−/y^* mutants (**Figure 1C**; p=0.0001). E-I ratio, calculated as E/(E+I) in each PYR cell, was increased in *Fmr1^−/y^* mice, demonstrating that IPCSs were reduced preferentially (p=0.0001). Note that E-I ratio is the ratio of synaptic conductances, measured with equal driving force on EPSCs and IPSCs. *Cntnap2^−/-^* mutants showed a similar phenotype relative to *Cntnap2^+/+^* littermates, with a somewhat more prominent reduction of inhibition (**Figure 1D**, n=12, 12 cells, p=0.0001). Data points for all individual cells are in **Figure S1**. Identical results were obtained when peak current amplitude was analyzed (**Figure S2**).

*16p11.2^del/+^* and *Tsc2^+/-^* mice showed qualitatively similar phenotypes, though more modest in magnitude (**Figure 1E-F**). IPSCs were reduced in both mutants (*16p11.2* vs. *16p11.2^+/+^*: n=15, 12 cells, p=0.002; *Tsc2* vs. Tsc2^+/+^: n=22, 15 cells, p=0.006), but feedforward EPSCs were not significantly reduced (*16p11.2*; p=0.36; *Tsc2*: p=0.17). This led to modestly increased E-I conductance ratio for both mutants (*16p11.2*: p=0.016, Tsc2: p=0.001). Results were identical for peak current amplitude (**Figure S2**). Overall, in *Fmr1^−/y^*, *Cntnap2^−/-^*, *16p11.2 ^del/+^* and *Tsc2^+/-^* mice, the area under the mean input-output curve for EPSCs was 0.57, 0.36, 0.86 and 0.92 of wild type respectively; for IPSCs it was 0.55, 0.18, 0.63 and 0.74 of wild type; and for E-I ratio was 2.24, 1.79, 1.29, 1.37 of wild type. Mutant and wild type PYR cells did not differ in baseline recording or stimulation parameters (**Table S2**), or in EPSC or IPSC kinetics including latency and EPSC-IPSC delay (**Table S2, Figure S3**). Thus, these 4 genetically distinct ASD mutants exhibited a common impairment in feedforward IPSCs, variably coupled to a loss of feedforward EPSCs, yielding a common increase in E-I conductance ratio.

Analysis of spontaneous miniature synaptic currents (mEPSCs and mIPSCs) in L2/3 PYR cells also revealed a preferential reduction in mIPSC activity compared to mEPSC activity, observed in 3 of the 4 ASD mutants (see Supplemental Information and **Figure S4**).

### Spiking excitability in the L2/3 network

Does increased E-I ratio drive stronger synaptic responses and more spiking in L2/3, as commonly predicted from the E-I ratio hypothesis? To test this, we first measured spontaneous spiking in L2/3 PYR neurons in slices bathed in low-divalent Active Ringers solution. Active Ringers is more similar in ionic composition to natural cerebrospinal fluid, and promotes spontaneous network activity in slices (Dani et al., 2005,Yassin et al., 2010). Cell-attached recording was used to preserve the normal intracellular milieu. Many L2/3 PYR cells showed spontaneous firing, which was abolished by APV and NBQX (100 μM and 10 μM; n=7 cells), indicating that it was driven by network synaptic activity (**Figure 2A-B**). We compared the distribution of L2/3 PYR firing rates in each ASD mutant genotype versus corresponding wild type (**Figure 2C**; n=45-66 cells per genotype). Surprisingly, *Fmr1^−/y^*, *16p11.2^del/+^* and *Tsc2^+/-^* mutants showed normal firing rates relative to wild type, and only *Cntnap2^−/-^* showed excess spiking (p = 0.033, KS test).

**Figure 2:**
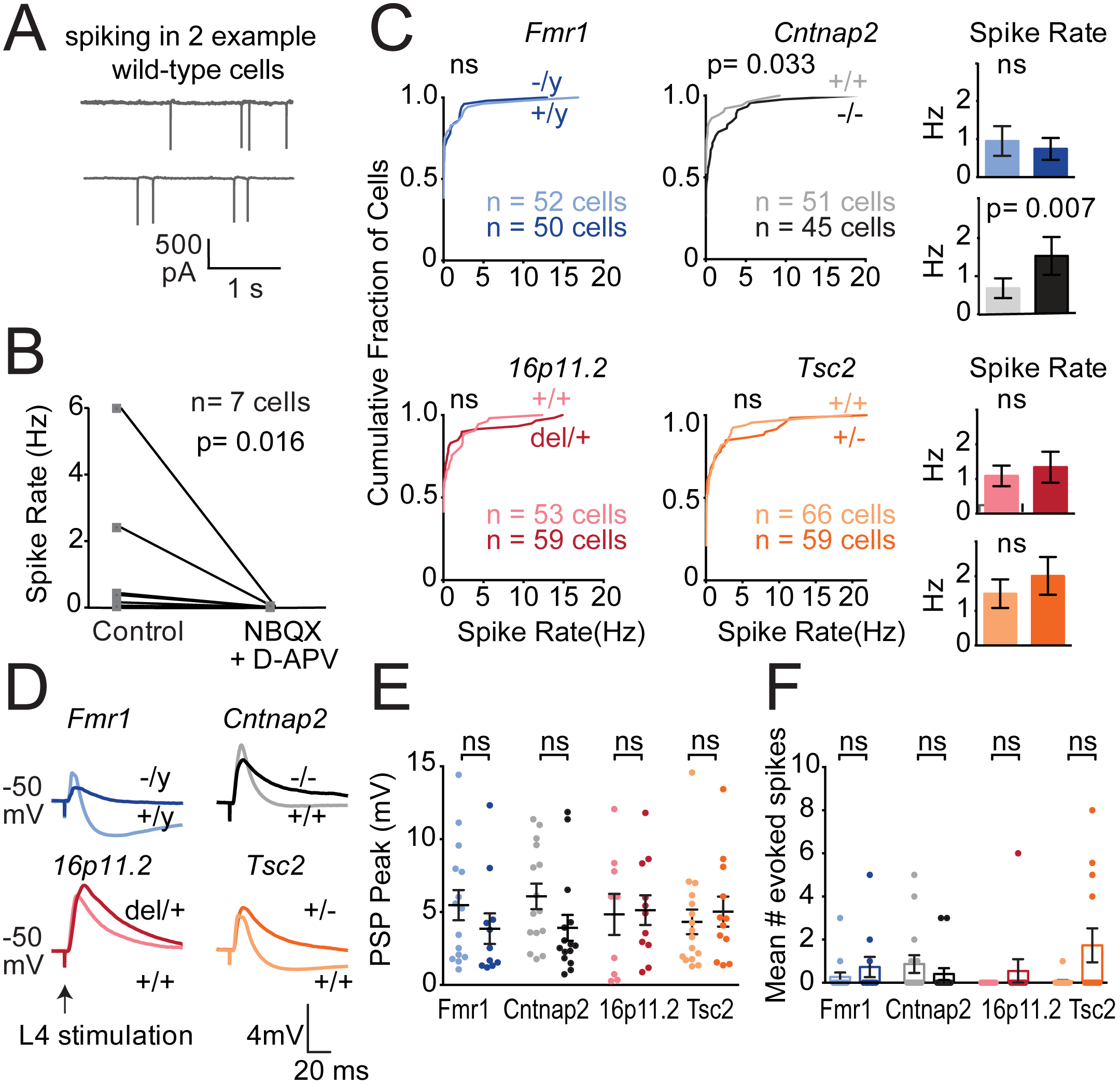
Spiking of L2/3 PYR cells in S1 slices. **(A-C)** Spontaneous spiking of L2/3 PYR cells in active slices. **(A)** Spontaneous spiking in active Ringer’s in two example L2/3 PYR cells in cell-attached mode. **(B)** Spontaneous spiking is abolished by glutamate blockers (n=7 L2/3 PYR cells from 4 wild type C57BL/6 mice). P-value from Wilcoxon matched-pairs signed rank test. **(C)** Cumulative distribution of spontaneous firing rate across all neurons in wild type (lighter colors) and ASD mutants (darker colors). Bars show mean ± SEM of the same data. Differences were assessed by KS test. **(D-F)** L4-evoked PSPs and spiking recorded in L2/3 PYR cells in current clamp. Recordings were from baseline Vm of −50 mV, with NMDA currents intact. **(D)** Mean waveforms of L4-evoked PSPs recorded in current-clamp mode from control and mutant L2/3 PYR neurons. Stimulus intensity was 1.4x Eθ for each cell.. **(E)** PSP peak amplitude for all cells (dots). Lighter colors, wild type. Darker colors, ASD mutants. Bars show mean ± SEM. n = 9-16 cells per genotype. **(F)** Mean number of L4-evoked spikes. Each dot is a cell. Bars show mean ± SEM. Differences in (E) and (F) were assessed by Mann-Whitney test, α= 0.05.

To understand why firing rate was largely normal in ASD mutants, we measured L4-evoked postsynaptic potentials (PSPs) and L4-evoked spiking in L2/3 PYR neurons. Recordings were made from a baseline Vm of −50 mV, i.e. in the just-subthreshold regime most relevant to natural, synaptically evoked spiking. For each cell, we first determined Eθ in voltage clamp, then switched to current clamp, depolarized the cell to −50±1.3 mV and measured L4-evoked PSPs and spikes at 1.4x Eθ. L4-evoked spiking was rare (5.2% of all sweeps, 17% of all cells), and PSPs were quantified from non-spiking sweeps. Example L4-evoked PSPs are shown in **Figure 2D**. Strikingly, no mutant genotype showed a PSP peak (maximum depolarization) greater than wild type (**Figure 2E**). Instead, PSP peak was unchanged from wild type (*16p11.2^+/+^* vs. *16p11.2^del/-^*: 4.8 ± 1.4 vs. 5.1 ±1.0 mV, n= 9, 11 cells, p=0.82, Mann-Whitney; *Tsc2^+/+^* vs. *Tsc2^+/-^*: 4.3±0.8 vs. 5.0±1.0 mV, n= 16, 12 cells, p=0.07) or showed a non-significant trend for weaker PSPs (*Fmr1^−/y^* vs. *Fmr1^−/y^*: 5.5±1.0 vs. 3.9 ±1.0 mV, n= 15, 11 cells, p=0.24; *Cntnap2^+/+^* vs. *Cntnap2^−/-^*: 6.1± 0.9 vs. 3.9± 0.9 mV, n= 15, 15 cells, p=0.07). The mean number of L4-evoked spikes was normal in ASD mutants, except for a modest trend toward more spikes in *Tsc2^+/-^* (*Fmr1*: p=0.49, *Cntnap2*: p=0.30, *16p11.2*: p=0.99, *Tsc2*: p=0.07, Mann-Whitney) (**Figure 2F**). There was also no difference in the fraction of cells that exhibited L4-evoked spiking (*Fmr1*: p=0.62, *Cntnap2*: p=0.39, *16p11.2*: p=0.99, *Tsc2*: p=0.14, Fisher’s exact test) (**Table S3**). Thus, despite the strong preferential loss of L4-evoked IPSCs in L2/3 PYR cells, L4-evoked synaptic responses and spiking were normal across ASD mutants, and spontaneous network spiking in active slices was only increased in *Cntnap2^−/-^* mice.

We also examined intrinsic excitability. L2/3 PYR cells showed normal passive properties at rest (**Table S4**). Intrinsic spiking excitability was variably affected across mutants, with no consistent phenotype (**Figure S5**; see Supplemental Information).

### Effects of increased E-I ratio evaluated using synaptic conductance model

To understand how reduced inhibitory conductance and increased E-I ratio could yield stable PSPs and evoked spiking, we modeled how L4-evoked excitatory and inhibitory synaptic conductances (G_ex_ and G_in_) generate PSPs in L2/3 PYR cells. For each neuron in Figure 1, we converted the EPSC and IPSC measured at 1.4x Eθ into G_ex_ and G_in_ waveforms, and then used a standard, passive parallel conductance model (Wehr and Zador, 2003) to predict the PSP that these conductances would elicit (**Figure 3A-B**). PSPs were modeled from a starting Vm of −50 mV to assess synaptic drive just below spike threshold. Model capacitance and resting conductance were from measured values for each genotype (**Table S4**). The model had no free parameters.

**Figure 3:**
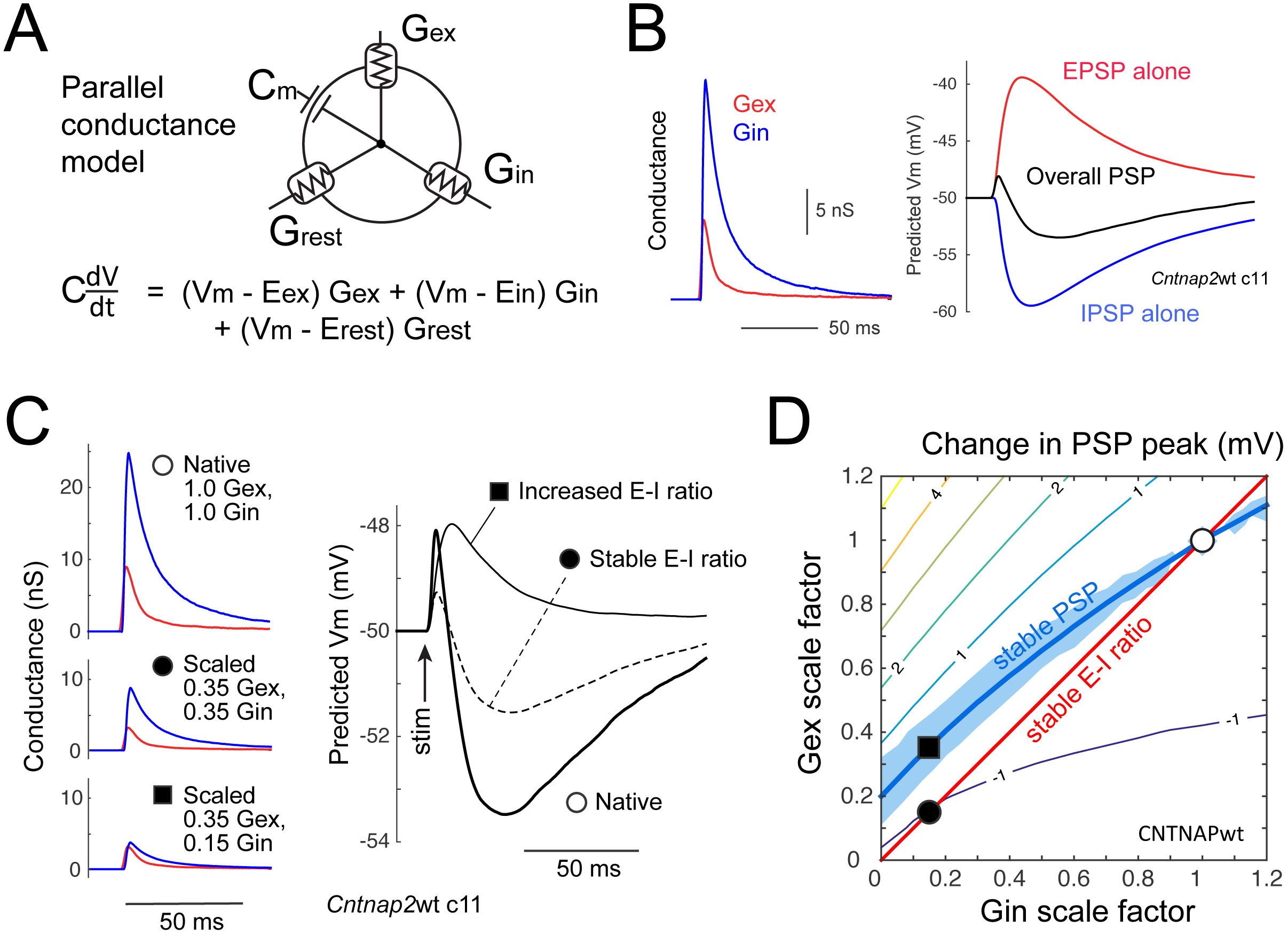
Relationship between E-I conductance ratio and PSP peak for cells near spike threshold. **(A)** Schematic of parallel conductance model. **(B)** Gex and Gin waveforms for an example wild type cell, and predicted EPSP (from Gex alone), IPSP (from Gin alone), and total PSP (from Gex and Gin together) at baseline V_m_ = −50 mV. **(C)** Conductance waveforms and predicted PSPs for one cell, for measured Gex and Gin waveforms at 1.4x Eθ (❍), after equal scaling to 0.35 of original (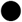), and further reduction in Gin to 0.15 of original that increases E-I conductance ratio (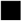). **(D)** Contour plot of mean predicted change in overall PSP peak for different combinations of Gex and Gin scaling, for all *Cntnap2^+/+^* cells. Thick contour shows Gex/Gin combinations that predict no change in PSP peak (PSP_diff_=0) from unscaled Gex/Gin. Blue region shows no significant change in PSP peak (p>0.05, bootstrap). Positive contour values denote increased predicted PSP peak. ❍ is average Gex and Gin in wild type cells. 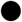 and 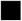 are from (C).

We first evaluated the standard claim that stable G_ex_:G_in_ ratio yields stable net synaptic depolarization (PSP peak), and that increasing G_ex_:G_in_ ratio increases PSP peak. Modeling showed this is incorrect. Instead, equal weakening of G_ex_ and G_in_ reduces PSP peak, and further weakening of G_in_ restores it (example cell, **Figure 3C**). The underlying principle is shown by a simulation in which we calculated the effect of differently scaled G_ex_ and G_in_ combinations on PSP peak for each *Cntnap2* wild type cell (**Figure 3D**). We predicted the PSP for each cell from its measured (unscaled) G_ex_ and G_in_ waveforms, and for combinations of G_ex_ and G_in_ scaled by factors of [0, 0.1, 0.2, … 1.2]. PSP peak for the unscaled G_ex_ and G_in_ combination was defined as PSP_unscaled_. PSP peak for all scaled G_ex_ and G_in_ combinations was expressed as PSP_diff_ = PSP_scaled_ – PSP_unscaled_. Averaging across wild-type cells revealed a smooth contour of G_ex_/G_in_ scaling combinations that predict no change in PSP peak (PSP_diff_ = 0), which we term the ‘PSP stability contour’ (**Figure 3D**, thick contour). This contour is above the diagonal when overall synaptic conductance weakens, indicating that G_in_ must decrease more than G_ex_ to maintain a constant PSP peak. This behavior is a result of lower driving force on G_in_ than G_ex_ for cells near spike threshold.

Next, we predicted PSPs from G_ex_ and G_in_ measured in ASD mutants. The mean EPSP peak predicted from G_ex_ alone was 2.0-4.3 mV smaller in ASD mutants than wild types (**Figure 4A**). This reduction was significant in *Cntnap2^−/-^*, *Fmr1^−/y^* and *Tsc2^+/-^* (2.4±0.4, 6.3±1.3, 13.3±2.1 mV) vs. wild type (6.8±1.2, 9.5±1.8, 15.9±1.7 mV, all p<0.037, KS test), but was only a trend in *16p11.2^del/+^* (9.0±1.2 vs. 11.0±2.0 mV for wild type). Similarly, the mean IPSP peak from G_in_ alone was predicted to be 1.9-4.3 mV lower in ASD mutants (**Figure 4A**). This was significant in *Cntnap2^−/-^*, *Fmr1^−/y^* and *Tsc2^+/-^* (1.1±0.4, 4.6±1.2, 8.6±1.1 mV) relative to wild types (5.4±1.0, 7.9±0.8, 11.9±0.8 mV, all p<0.024 KS test), but was a trend in *16p11.2^del/+^* (4.5±1.0 vs. 6.4±1.2 mV, p=0.19). Thus, reduced EPSCs and IPSCs in autism mutants predict smaller EPSPs and IPSPs near spike threshold. Combined G_ex_ and G_in_ waveforms generally predicted EPSP-IPSP sequences (**Figure 4B**). Peak of this overall PSP was identical between autism genotypes (*Cntnap2^−/-^* 1.6±0.4 mV, *Fmr1^−/y^* 1.9±0.3, *16p11.2^del/+^* 4.1±0.7, *Tsc2^+/-^* 3.7±0.8) and wild types (*Cntnap2^+/+^*1.5±0.3, *Fmr1^+/y^* 2.1±0.6, *16p11.2^+/+^* 3.6±0.5, *Tsc2^+/+^* 2.2±0.4 mV, all p>0.1, KS test). Across genotypes, the average difference in PSP peak was only 0.5 mV, even though the late IPSP was generally reduced (**Figure 4A-C**). Thus, EPSP and IPSP reductions counteract each other to stabilize PSP peak. To test this idea more thoroughly, we determined the PSP stability contour at 1.4x Eθ for wild types of each strain. We then plotted the mean change in G_ex_ and G_in_ magnitude observed in mutants at 1.4x Eθ (values from Figure 1, plotted as filled circles in **Figure 4D**). These points fell on, or within 0.5 mV of, the PSP stability contour from wild types. Thus, the reductions in G_ex_ and G_in_ in autism mutants are quantitatively matched to preserve synaptically-evoked peak ΔVm, not to increase it.

**Figure 4:**
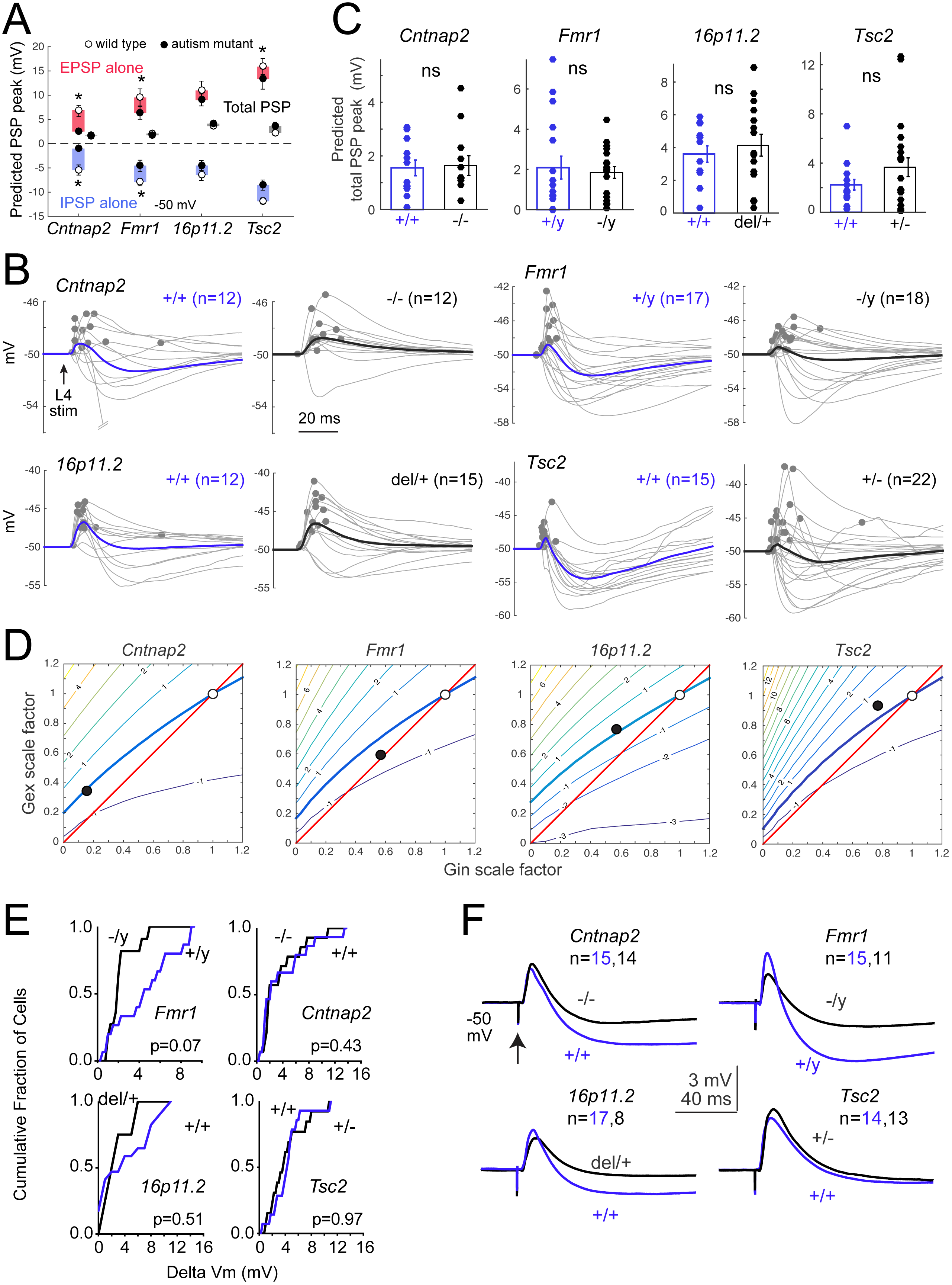
E-I conductance changes in ASD mutants predict stable PSPs. **(A)** Mean predicted EPSP, IPSP, and total PSP peak for each genotype at baseline Vm = −50 mV, for Gex and Gin recorded at 1.4x Eθ. Symbols are mean ± SEM across cells. N for each genotype is in (C). Stars, p<0.05, KS test. **(B)** PSP waveforms predicted from the measured Gex and Gin in each wild type (left) and mutant (right) cell. Dots show peak for each cell. Bold, mean predicted PSP across cells. **(C)** Distribution of peak PSP for each genotype. Bars show mean ± SEM. ns, not significant by KS test. **(D)** Contour plots show PSP stability contour (thick curve) for all wild type cells of each genotype. ❍, average Gex and Gin of wild type cells [(1,1) by definition]. 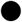, average Gex and Gin measured in mutant cells, as fraction of wild type. In all mutants, this lies within 0.5 mV of the PSP stability contour. **(E)** Cumulative histograms of measured L4-evoked PSP peak across cells in each genotype from baseline Vm of −50 mV, at 1.4x Eθ, with APV in bath. There were no significant differences in these distributions between any ASD mutant and its wild type. Statistics are by KS test, α=0.05. **(F)** Mean PSP waveforms for the experiment in (E).

We validated model predictions by recording L4-evoked PSPs in L2/3 PYR cells from −50 mV baseline Vm, this time with APV present to match conditions in the parallel conductance model, which lacks voltage-dependent NMDA currents. Stimulation was at 1.4x Eθ. Results were identical to the model predictions: PSP peak was unaffected, though the late IPSP was reduced in all mutants (**Figure 4E-F**; **Figure S6**). The only exception was a moderate but non-significant trend toward reduced PSP peak in *Fmr1^−/y^*, replicating the model results (**Figure 4E-F**).

To extend these predictions across the full physiological range of baseline Vm, we also modeled PSPs elicited from −70 mV. This model predicted weaker overall PSPs in mutants relative to wild type for *Cntnap2* and *Fmr1* (p<0.027, KS test), but not *16p11.2* and *Tsc2*. This is expected, because low driving force on inhibition at V_rest_ means that PSPs will track G_ex_, which is reduced in *Cntnap2* and *Fmr1* (**Figure S7**). Overall, the observed increase in E-I conductance ratio in these 4 ASD mutants predicts stable PSP amplitude for cells near spike threshold, and reduced PSP amplitude near V_rest_. The only deviation from this prediction was a non-significant trend toward reduced, not increased, PSP amplitude in *Fmr1^−/y^* mice near spike threshold (**Figure 4E-F**).

### L2/3 network activity and sensory coding in vivo

The results above suggest that despite substantial loss of inhibition, L2/3 spike rate may be relatively unchanged or even reduced *in vivo*. To test this, we recorded single units with laminar polytrodes in L4 and L2/3 of S1 in adult urethane-anesthetized mice (P42-92, mean P62), and measured spiking in response to calibrated whisker deflections. We tested *Cntnap2^−/-^*, *Fmr1^−/y^* and *16p11^del/+^* mice and corresponding wild types (**Figure 5**). Recordings were made in C1-2 and D1-2 whisker columns, identified by post-hoc histological staining or multiunit tuning in L4. We interleaved deflections of 9 single whiskers to map whisker receptive fields, plus deflections of the columnar whisker at multiple velocities to measure a velocity response curve (VRC) that parameterizes the gain and sensitivity of whisker-evoked spiking (**Figure 5A-B**). A median of 7 well-isolated single units were recorded in each animal. Individual units were classified as fast-spiking (FS; putative PV interneurons) or regular-spiking (RS; putative excitatory neurons) using a spike width criterion. This criterion was validated in separate experiments in which we recorded with the same electrodes in PV-Cre::ChR2 mice, and optogenetically identified spike waveforms of PV neurons from short-latency responses to blue laser flashes (**Figure 5C**).

**Figure 5:**
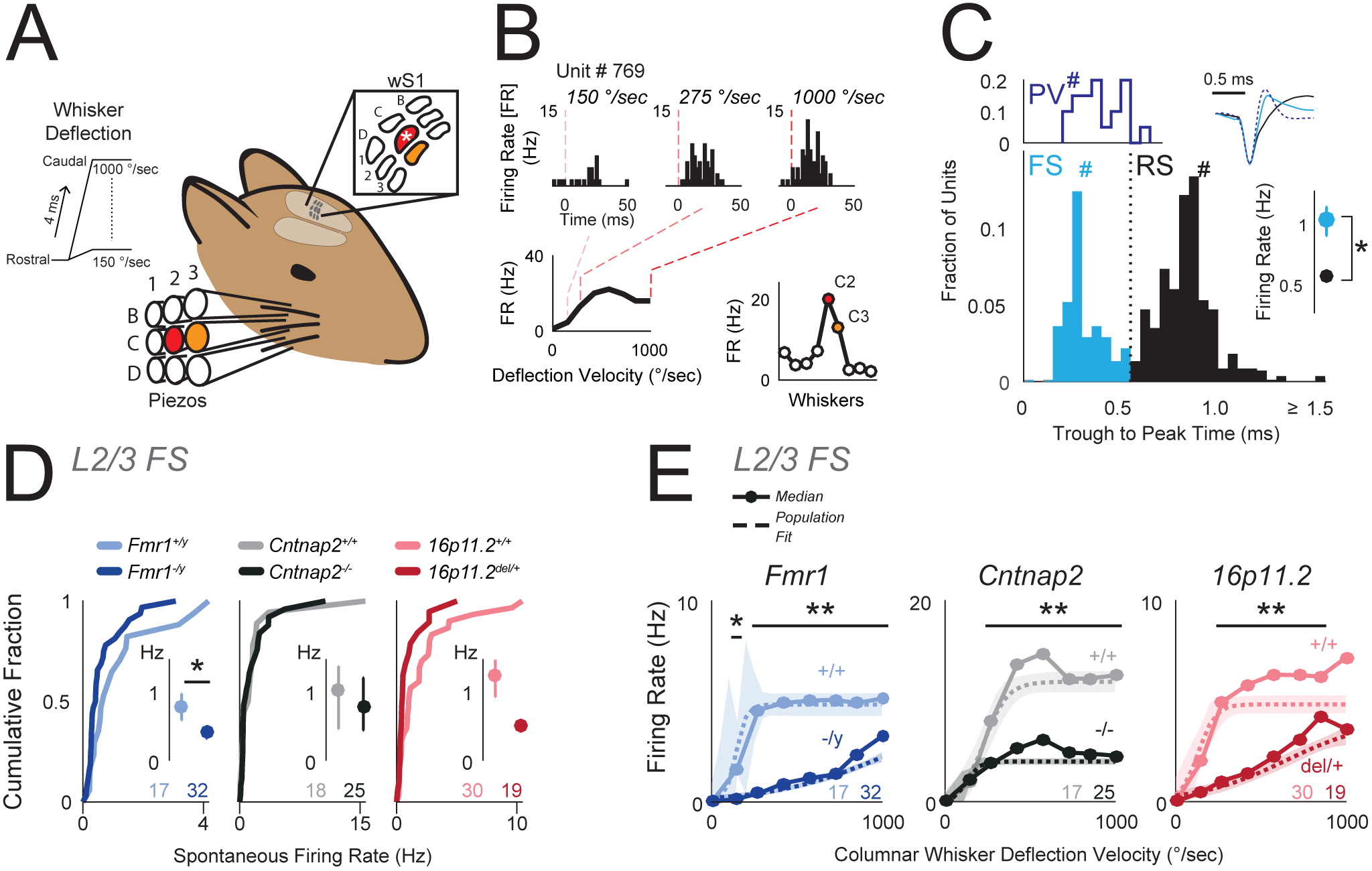
Reduced columnar whisker-evoked firing of inhibitory FS units in layer 2/3. **(A)** Schematic for in vivo recording experiments. 9 whiskers were individually deflected by a 3×3 array of piezoelectric actuators centered on the columnar whisker. Asterisk, recorded column. **(B)** Example L2/3 unit showing responses to columnar whisker deflection at 150, 275, and 1000 °/sec. Bottom, Velocity response curve (VRC; left) and whisker tuning curve (right) for this unit. Filled symbols, significant whisker response. **(C)** Top left: Trough-to-peak times for optogenetically-tagged PV neurons in PV-Cre::ChR2 mice. Bottom: Trough-to-peak times for units from ASD mutant and wild type mice. Dotted line, FS-RS threshold. Hashes mark the example waveforms (upper right). Right: Bootstrapped median firing rate for FS and RS units. Error bars are 68% CI. n = FS: [285, 69] (units, mice), RS: [546,69]. * p < 0.001, permutation test. **(D)** Spontaneous firing rate distributions for L2/3 FS units. Insets: Bootstrapped medians. Error bars are 68% CI. Numbers are units per genotype. * p = 0.04, permutation test. **(E)** Velocity response curves for the L2/3 FS unit population, calculated after subtraction of spontaneous rate for each unit. Circles and solid lines: Population median firing rate. Dashed curve is sigmoid fit to population data. Shaded region is 68% CI. Numbers are units per genotype. * p = 0.0066, ** p << 0.0001, t-test.

We first tested whether reduced inhibition in L2/3 of ASD mutants was reflected in FS unit spiking. Spontaneous spiking of L2/3 FS units was significantly reduced in *Fmr1^−/y^* mice and showed non-significant trends toward reduced rate in the other ASD mutants (**Figure 5D**) (bootstrapped median firing rate [Hz]: *Fmr1^+/y^* 0.76, *Fmr1^−/y^* 0.40, p = 0.04; *Cntnap2^+/+^* 0.99, *Cntnap2^−/-^* 0.77, p = 0.83; *16p11.2^+/+^* 1.20, *16p11.2^del/+^* 0.50, p = 0.08, permutation test). N’s for *in vivo* measurements are in **Table 5**. Whisker-evoked spiking of L2/3 FS units was measured in the VRC, which reflects feedforward activation of FS inhibitory circuits. For each genotype, population VRC data was fit with a sigmoid to quantify response threshold (the deflection velocity that evokes half-maximal response), sensitivity and maximal evoked firing rate (**Figure 5E**). All three ASD mutant genotypes showed significant decreases in maximal whisker-evoked firing rate for L2/3 FS units (**Figure 5E**, dashed lines, p < 0.007, permutation test). This was also apparent in the median response across recorded units (**Figure 5E**, solid lines), and in a reduction in total whisker-evoked spikes across all velocities (median spike count: *Fmr1^+/y^* 36.03, *Fmr1^−/y^* 11.92, p = 0.002; *Cntnap2^+/+^* 102.72, *Cntnap2^−/-^* 33.04, p = 0.126; *16p11.2^+/+^* 43.52, *16p11.2^del/+^* 19.13, p = 0.05, permutation test). Response thresholds were not altered, except for a modest decrease in *Cntnap2^−/-^* mice (p = 0.047, t-test). This common reduction in whisker-evoked spiking of L2/3 FS neurons suggests that feedforward inhibition is reduced *in vivo*, as in S1 slices.

To test whether L2/3 PYR activity was abnormal, we analyzed spiking of L2/3 RS units. Spontaneous spiking of L2/3 RS units was normal in ASD mutants relative to wild types (**Figure 6A**; bootstrapped median [Hz]: *Fmr1^+/y^* 0.58, *Fmr1^−/y^* 0.32, p = 0.055; *Cntnap2^+/+^* 0.30, *Cntnap2^−/-^* 0.41, p = 0.17; *16p11.2^+/+^* 0.50, *16p11.2^del/+^*0.61, p = 0.85, permutation test). The fraction of L2/3 RS units that were whisker-responsive was also normal (*Fmr1^+/y^* 0.39, *Fmr1^−/y^* 0.53, p = 0.10; *Cntnap2^+/+^* 0.26, *Cntnap2^−/-^* 0.39, p = 0.24; *16p11.2^+/+^* 0.46, *16p11.2^del/+^* 0.59, p = 0.19, χ^2^ test) (**Figure S8A**). Whisker-evoked spiking across all units in the VRC was normal in *Cntnap2^−/-^* and *16p11.2^del/+^* mice, and was actually reduced in *Fmr1^−/y^* mice relative to wild type (**Figure 6B**, p < 0.003, t-test). VRC response threshold was also unchanged in ASD mutants (data not shown). The total number of whisker-evoked spikes across the VRC was not altered in any ASD mutant (median spike count: *Fmr1^+/y^* 20.14, *Fmr1^−/y^* 10.26, p = 0.12; *Cntnap2^+/+^* 11.27, *Cntnap2^−/-^* 12.09, p = 0.16; *16p11.2^+/+^* 25.44, *16p11.2^del/+^* 30.145, p = 0.85, permutation test). Finally, the mean spiking response to each unit’s preferred (best) whisker was also normal (**Figure S8B**). Thus, whisker-evoked population firing rate in L2/3 RS cells was normal, not elevated, in *Cntnap2^−/-^* and *16p11.2^del/+^* mice, and was reduced in *Fmr1^−/y^* mice, despite strongly reduced inhibition in these genotypes.

**Figure 6:**
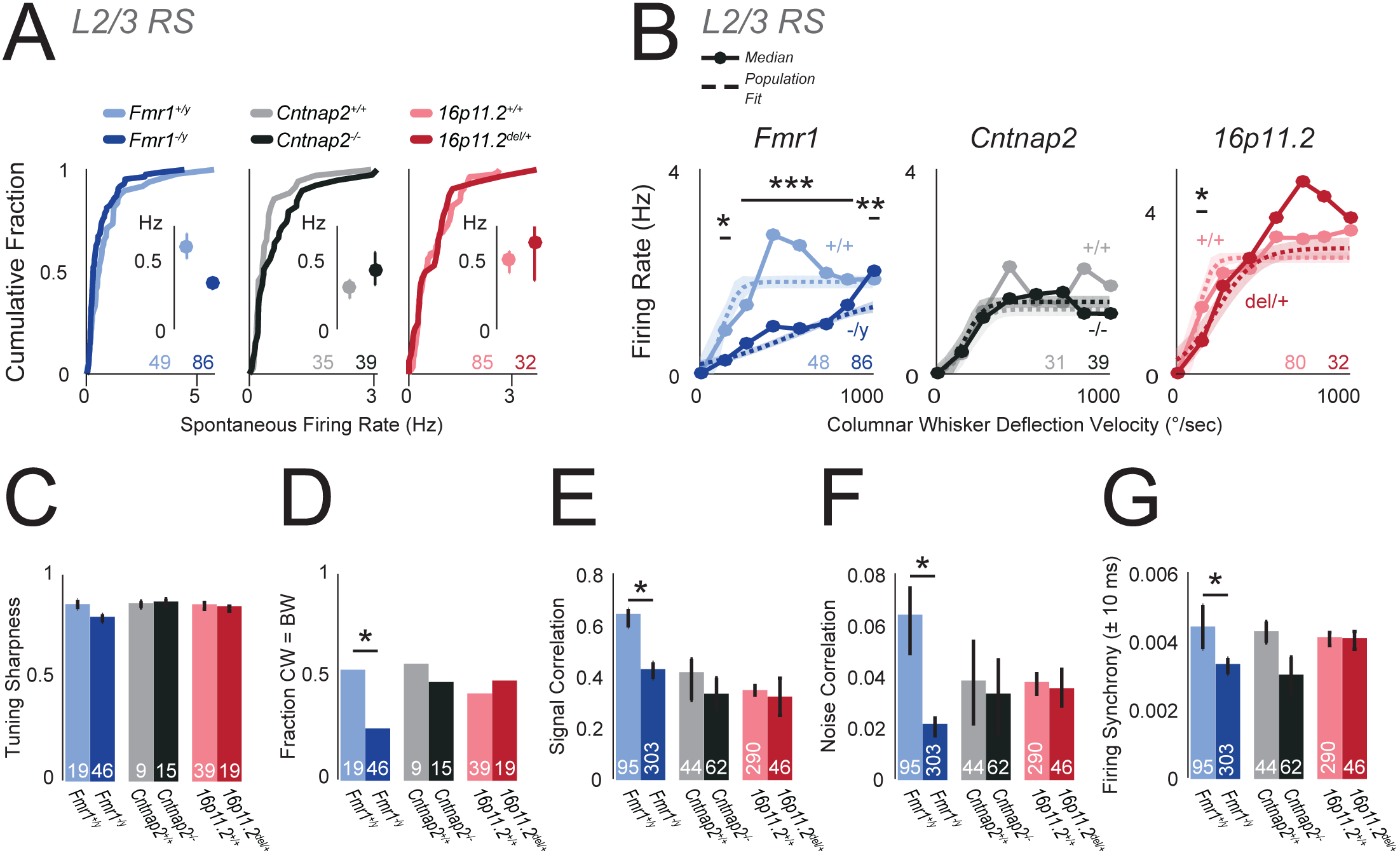
Firing of excitatory L2/3 RS units *in vivo* is largely normal in autism mutants. **(A)** Spontaneous firing rate distributions for L2/3 RS units. Insets: Bootstrapped medians. Error bars, 68% CI. In all panels, number of units per genotype are shown. **(B)** Velocity response curves for the L2/3 RS unit population, calculated after subtraction of spontaneous rate for each unit. Circles and solid lines: Population median. Dashed curve is sigmoid fit to population data. Shaded region is 68% CI. Numbers are units per genotype. * p < 0.003, ** p = 0.0003, *** p << 0.0001, t-test. **(C)** Tuning sharpness of whisker-responsive units. Bars, bootstrapped median. Error bars, 68% CI. **(D)** Fraction of whisker-responsive units whose best whisker (BW) is the columnar whisker (CW). * p = 0.0243, χ^2^ test. **(E)** Signal correlation for pairs of L2/3 RS neurons. Bars, bootstrapped median. Error bars: 68% CI. * p << 0.0001, permutation test. **(F)** Noise correlation for pairs of L2/3 RS neurons. Bars, bootstrapped median. Error bars: 68% CI. * p = 0.0005, permutation test. **(G)** Raw firing synchrony for pairs of L2/3 RS neurons, calculated as mean over ±10 ms in the cross-correlogram. Bars, bootstrapped median. Error bars: 68% CI. * p = 0.00027, permutation test.

### Sensory tuning and firing correlations

Inhibition regulates spike timing and sensory tuning in sensory cortex, in addition to firing rate (Gabernet et al., 2005,Wehr and Zador, 2003). We tested whether L2/3 RS units showed deficits in any of these sensory coding properties, which could add noise to circuits. We found no deficits in spike latency or jitter, except for a modest reduction in spike latency in *16p11.2^del/+^* mice (**Figure S8C-D**). Tuning sharpness was not impaired in any ASD genotype (**Figure 6C**). *Fmr1^−/y^* mice show spatially broader cortical activation to single-whisker stimulation, implying a blurred, less organized whisker map (Juczewski et al., 2016,Zhang et al., 2014). Consistent with a blurred map, we found that the fraction of L2/3 RS units that were tuned to the columnar whisker was lower in *Fmr1^−/y^* mice (*Fmr1^+/y^* 0.53, *Fmr1^−/y^* 0.24, p = 0.024, χ^2^ test) (**Figure 6D**). This effect was also observed in *Fmr1^−/y^* as a decrease in pairwise tuning similarity (signal correlation) between simultaneously recorded L2/3 RS neurons (**Figure 6E**). However, neither *Cntnap2^−/-^* nor *16p11.2^del/+^* mutants shared these phenotypes (**Figure D-E)**. Thus, sensory tuning was remarkably normal in ASD mutants, except for a blurring of the whisker map in *Fmr1^−/y^* animals.

Inhibition also regulates local cortical rhythms and firing correlations, which can strongly impact population coding. We calculated trial-by-trial spike count correlations (noise correlations) for pairs of simultaneously recorded L2/3 PYR cells (median 6 pairs with < 200 μm inter-cell distance per mouse) as well as raw firing synchrony, calculated as mean correlation at 0±10 ms time lag from the spike cross-correlogram. *Fmr1^−/y^* mice showed significantly reduced noise correlations relative to *Fmr1^+/y^* controls, but *Cntnap2^−/-^* and *16p11.2^del/+^* showed no change (**Figure 6F**). *Fmr1^−/y^* and *Cntnap2^−/-^* mice showed a similar tendency for reduced firing synchrony vs. wild types, but this was significant only for *Fmr1^−/y^* mice (*Fmr1^+/y^* vs *Fmr1^−/y^,* p = 0.00027; *Cntnap2^+/+^* vs *Cntnap2^−/-^,* p = 0.07; *16p11.2^+/+^* vs *16p11.2^del/+^*, p = 0.49, permutation test) (**Figure 6G**). Thus, firing correlations were decreased or unchanged in ASD mutants.

### Sensory-evoked spiking in L4

ASD mutants showed generally normal L4-L2/3 feedforward PSPs *in vitro*, with a trend toward reduced PSP peak in *Fmr1^−/y^* mice (**Figures 2**,**4**). Sensory gain between L4 and L2/3 *in vivo* may parallel these *in vitro* measurements of functional synaptic strength. To test this, we measured spiking of L4 RS units, typically recorded after L2/3 in the same penetrations. Spontaneous activity of L4 RS units was normal across all ASD mutant genotypes (**Figure S9A**). Whisker-evoked spiking in the VRC for L4 RS units was normal for *Fmr1^−/y^* and *16p11.2^del/+^* mice (**Figure S9B**). This suggests that the effective sensory gain between L4 and L2/3 was reduced in *Fmr1^−/y^*, and was normal in *16p11.2^del/+^*, matching the L4-L2/3 synaptic phenotypes in these mutants. In contrast, *Cntnap2^−/-^* L4 RS units had abnormally low whisker-evoked spiking (**Figure S9B**, p < 0.007, t-test). Thus, the existence of normal whisker-evoked spiking in L2/3 in this mutant suggests that sensory gain between L4 and L2/3 was increased in *Cntnap2^−/-^*, perhaps related to the increased network excitability observed in active slices (**Figure 2**).

## Discussion

### Common increase in E-I conductance ratio

Despite its prominence, systematic tests of the E-I ratio hypothesis across different genetic forms of ASD are lacking. We provide the first broad test of this hypothesis at the synapse and circuit physiology levels, in 4 genetically distinct ASD mouse models. We found a common phenotype of decreased L4-L2/3 feedforward inhibition and a smaller, variable decrease in feedforward excitation, yielding a common decrease in total synaptic conductance and increase in E-I conductance ratio in L2/3 PYR cells. mIPSCs were generally reduced more than mEPSCs in ASD mutants, suggesting a broad circuit phenotype of reduced inhibition. *MeCP2^−/y^* mice exhibit a qualitatively similar combination of strongly reduced inhibition and more modestly reduced excitation in L2/3 of visual cortex (Banerjee et al., 2016), and *Ube3a^m-/p+^* have a similar phenotype (Wallace et al., 2012). Thus, at least 5, and possibly 6 well-validated ASD mouse models share a similar loss of total synaptic conductance, loss of inhibition and increase in E-I conductance ratio in L2/3 of sensory cortex.

These results extend prior findings of reduced inhibition in *Fmr1^−/y^* mice from L4 (Gibson et al., 2008) to L2/3, and in *Cntnap2^−/-^* from hippocampus (Jurgensen and Castillo, 2015) to neocortex. It is also consistent with reduced inhibitory neuron number and PV expression in *Fmr1^−/y^* and *Cntnap2^−/-^* (Peñagarikano et al., 2011,Selby et al., 2007). *Tsc2^+/-^* and *16p11.2^del/+^* phenotypes were previously unknown. Overall, reduced inhibition and increased E-I ratio appear to be more common in sensory cortex than defects in mGluR-related protein synthesis and synaptic plasticity, which have opposite effects in *Tsc2^+/-^* and *Fmr1^−/y^* mice (Auerbach et al., 2011).

### Network spiking excitability is largely preserved

Despite elevated E-I ratio, excess spiking did not occur among L2/3 PYR cells, contrary to the standard E-I ratio hypothesis. In standard slice conditions, L4-evoked spiking in L2/3 PYR cells from just-subthreshold Vm was normal in all mutants, as was peak depolarization during L4-evoked PSPs. In active slices, 3 of 4 mutant genotypes had normal spontaneous firing. *In vivo*, all 3 ASD mutants tested showed reduced whisker-evoked spiking of L2/3 FS units, consistent with reduced feedforward inhibition. However, spiking of L2/3 RS (presumed excitatory) units was normal in *Cntnap2^−/-^* and *16p11.2^del/+^* mice, and was reduced in *Fmr1^−/y^* mice. Thus, increased E-I ratio in the L4-L2/3 projection was associated with remarkably normal evoked synaptic responses and spiking in L2/3 PYR cells, and even with reduced firing in *Fmr1^−/y^ in vivo*. Only *Cntnap2^−/-^* mice showed hints of increased spiking excitability in L2/3, apparent from spontaneous spiking in active slices and in increased response gain from L4 to L2/3 *in vivo*.

Many prior studies of *in vivo* spiking activity in ASD mutants also show normal or reduced cortical firing rates. Spontaneous firing rate is normal in L2/3 of S1 and V1 in *Fmr1^−/y^*, *Cntnap2^−/-^*, *Pten^−/^*^−^ and *Ube3a^m-/p+^* mice (Garcia-Junco-Clemente et al., 2013, O’Donnell et al., 2017, Peñagarikano et al., 2011,Wallace et al., 2017), and reduced in V1 of *MeCP2^−/y^* mice (Durand et al., 2012). Sensory-evoked spike rate and population activity are normal in L2-4 of S1 and V1 in *Fmr1^−/y^* and *Ube3a^m-/p+^* (Dolen et al., 2007, Juczewski et al., 2016,Wallace et al., 2017), reduced in L2/3 of V1 in *MeCP2^−/y^* and *Pten^−/-^* (Banerjee et al., 2016, Durand et al., 2012,Garcia-Junco-Clemente et al. 2013) and slightly reduced in S1 in *Nlgn4^−/-^* mice (Unichenko et al., 2017). Increased sensory-evoked spiking has only been observed in a small sample of hindpaw S1 neurons (Zhang et al., 2014) and as a modest increase in late spikes in auditory cortex (Rotschafer and Razak, 2013), both in *Fmr1^−/y^*. Thus, despite the prevalence of seizures in some ASD genotypes, increased cortical spiking is generally not observed. Increased network excitability is instead suggested by subtler phenotypes, including modestly elevated firing correlations and longer UP states in young *Fmr1^−/y^* mice (Goncalves 2013, Hays et al., 2011, O’Donnell et al., 2017), increased intra-burst spike frequency in *Shank3B^−/-^* mice (Peixoto et al., 2016), and broader sensory tuning in *MeCP2^−/y^*, *Pten^−/-^, Fmr1^−/y^* and *Ube3a^m-/p+^* mice (Banerjee et al., 2016, Garcia-Junco-Clemente et al. 2013, Juczewski et al., 2016,Wallace et al., 2017). *Fmr1^−/y^* mice show faster or further spread of sensory-evoked activity in S1, suggesting a blurred whisker map (Arnett et al., 2014; Zhang et al., 2014). We also observed whisker map blurring in *Fmr1^−/y^* mice, in the form of increased tuning heterogeneity in each S1 column.

### E-I ratio is coordinated to stabilize synaptic responses near spike threshold

A simple synaptic conductance model explains why increased E-I conductance ratio does not generate stronger PSPs or more spiking in ASD mutants: In all 4 ASD genotypes, the decreases in inhibitory and excitatory conductances were precisely balanced to maintain stable PSPs, for Vm just below spike threshold. This Vm range is most relevant for naturally evoked spiking, as observed during active touch sequences *in vivo* (Yamashita et al., 2013). Because driving force is less for inhibition than excitation in this Vm range, the relatively large decrease in feedforward G_in_ (to 0.15-0.57 of wild type, for the 4 ASD genotypes) and the smaller decrease in G_ex_ (to 0.35-0.92 of wild type) predict equal, opposing reductions in IPSP and EPSP amplitude. Together, these preserve PSP peak in all 4 ASD mutants (**Figure 4**). Simulations defined a smooth contour of G_in_ and G_ex_ reductions that jointly stabilize feedforward PSP peak, for just-subthreshold baseline Vm (**Figure 4D**). The mean G_in_ and G_ex_ reduction was close to this PSP stability contour in all 4 ASD mutants, and predicted < 0.5 mV change in PSP peak. Measurement of L4-evoked PSPs and spikes in L2/3 PYR cells confirmed that neither PSPs nor spikes were significantly altered in ASD mutants, despite the pronounced reduction in G_ex_ and G_in_ (**Figure 4**).

Thus, the common interpretation that increased E-I synaptic conductance ratio necessarily predicts increased spiking excitability in networks is incorrect. Instead, the specific increase in E-I conductance ratio offsets the decrease in total synaptic conductance in these 4 ASD genotypes to produce stable PSPs. Stable PSPs may also occur in *MeCP2^−/y^* mice, where visual-evoked G_ex_ and G_in_ are reduced to ~0.60 and ~0.45 of wild type in L2/3 PYR cells (Banerjee et al., 2016), which is numerically similar to the 4 ASD mutants tested here. The simulation also predicts stable PSP peak when if Gex and Gin both increase while E-I ratio decreases (**Figure 4D**), as in one ASD model (Harrington et al., 2016). Thus, functionally matched changes in Gex and Gin that alter E-I ratio but preserve PSP peak are a common theme across a diverse set of ASD genotypes.

These predictions do not account for active conductances including NMDA receptors, shunting inhibition, or changes in GABA_A_ reversal potential which occur in young *Fmr1^−/y^*, *MeCP2^−/y^*, and valproate models of ASD (Banerjee et al., 2016, He et al., 2014,Tyzio et al., 2014). Despite this, these predictions explain the largely stable firing rate in S1 *in vivo* and in active slices for 3 of 4 ASD mutants. Interestingly, *Fmr1^−/y^* was the only genotype to show a trend for weaker feedforward PSPs *in vitro* (**Figures 2**, **4**), and this mouse also showed reduced whisker-evoked spiking in L2/3 *in vivo* (**Figure 6**). In all ASD mutants, while the PSP peak remained stable, the late IPSP following the peak was substantially weakened (**Figure 4**). This suggests that sparsely active cortical areas like L2/3 of S1 which receive discrete, low-frequency volleys of synaptic input (Barth and Poulet, 2012) may exhibit stable spiking due to preservation of PSP peak, but other areas that receive denser input could exhibit excess spiking due to enhanced temporal summation during input trains.

### E-I ratio and synaptic homeostasis in autism

Our results show that increased E-I conductance ratio is common across ASD genotypes, but yields stable synaptic drive and largely stable spiking, at least in L2/3 of sensory cortex. How then is elevated E-I ratio related to information processing deficits in ASD? Our results strongly suggest that E-I ratio changes are compensatory in autism (Nelson and Valakh, 2015). Both excitatory and inhibitory circuits exhibit robust homeostatic plasticity that adjusts E-I ratio to stabilize cortical firing rate (Gainey and Feldman, 2017,Turrigiano 2011). In S1, this E-I homeostasis is evident during brief whisker deprivation, which weakens L4-L2/3 inhibition more than excitation, increasing E-I ratio by a precise amount that maintains stable PSPs and spiking in L2/3 (Gainey et al., 2018,Li et al., 2014). This is virtually identical to the phenotype in ASD mutants (**Figure S10**). We propose that ASD mutations alter cortical spiking activity, which secondarily engages E-I homeostasis to restore cortical firing rate. ASD symptoms may arise from imperfect homeostasis that largely normalizes firing rate but compromises subtler aspects of population coding, like firing synchrony. Alternatively, elevated E-I ratio may impair the capacity to compensate for future challenges or strong inputs (Ramocki and Zoghbi, 2008), as in audiogenic seizures (Rotschafer and Razak, 2013).

This compensatory model explains why diverse genetic mutations all alter E-I ratio, why firing rate is only modestly affected, and why G_ex_ and G_in_ changes are coordinated to stabilize PSPs. Because E-I homeostasis is a natural response to network perturbation, E-I ratio changes are expected in numerous neurological disorders, as has been observed (Selten et al., 2018). This view predicts that enhancing inhibition may be insufficient to normalize ASD symptoms in cases or brain areas where effective E-I homeostasis (i.e., that normalizes cortical spike rate) has taken place.

## Author Contributions

MA, PS, TL: Designed, performed and analyzed in vitro (MA) and in vivo (PS, TL) experiments. DF: Designed experiments and performed modeling. MA, TL, DF: Co-wrote paper. DF: Secured funding.

## Acknowledgments

We thank Michael Nemeh, Michelle Ju, Theodore Huynh for technical assistance, and Melanie Gainey for advice. This work was supported by the Simons Foundation (342096), the Miller Institute for Basic Research at UC Berkeley (MA), and the Ford Foundation (MA).

## Methods

ASD model mice were obtained from Jackson Labs (Fmr1^+/y^: #004828, Fmr1^−/y^: #004624; Cntnap2^+/-^: #017482; 16p11.2 ^del/+^: #013128; Tsc2^+/-^: #004686). Genotyping was by PCR, using Jackson Lab protocols. Optogenetic tagging experiments were performed in PV-Cre^mut/+^::ChR2^mut/+^ mice, bred by crossing PV-Cre JAX #017320 with Ai32 JAX #024109. Mice were maintained on a 12:12-hr light-dark cycle. Mice were group-housed, and weaned at postnatal day (P) 21. Slice physiology experiments used male mice aged P17-P23. *In vivo* physiology experiments used male mice aged P42-P92. All procedures were approved by the Institutional Animal Care and Use Committee at UC Berkeley.

## Slice Preparation

S1 slices (350 μm thick) from P17-23 mice were cut using standard methods in the “across-row” plane oriented 35° toward coronal from midsagittal, which allows unambiguous identification of whisker barrel columns (House et al., 2011).. Cutting solution contained (in mM): 85 NaCl, 75 sucrose, 25 D- (+)-glucose, 4 MgSO_4_, 2.5 KCl, 1.25 NaH_2_PO_4_, 0.5 ascorbic acid, 25 NaHCO_3_, 0.5 CaCl_2_. Slices were then incubated at 32°C for 30 min in standard Ringer’s solution (in mM: 119 NaCl, 2.5 KCl, 1.3 MgSO_4_, 1 NaH_2_PO_4_, 26.2 NaHCO_3_, 11 D-(+)-glucose and 2.5 CaCl_2_; all solutions were pH 7.3, 300 mOsm, and bubbled with 95% O_2_ and 5% CO_2_). Slices were maintained at room temperature >30 min before recording.

## *In Vitro* Physiology

Synaptic physiology recordings were made at 30°C in standard Ringer’s solution (2.5-3.0 mL/min). Spontaneous spiking was recorded at 35°C in active Ringer’s solution, which was identical to standard Ringer’s except that it contained 3.5 mM KCl, 0.25 mM MgSO_4_ and 1mM CaCl_2_.

Whole-cell recordings were targeted using infrared DIC optics. L2/3 PYR cells were identified visually, and regular spiking was verified in current clamp. Recordings were made using 3–6 MΩ pipettes and a Multiclamp 700B amplifier (Molecular Devices, Sunnyvale, CA). Signals were low-pass filtered (2-6 kHz) and digitized (10-20 kHz).

For voltage clamp recordings, the internal contained (in mM): 108 D-gluconic acid, 108 CsOH, 20 HEPES, 5 tetraethylammonium-Cl, 2.8 NaCl, 0.4 EGTA, 4 MgATP, 0.3 NaGTP, 5 BAPTA, 5 QX314 bromide (adjusted to pH 7.2 with CsOH, 290 mOsm). For current clamp recordings, the internal contained: 116 K gluconate, 20 HEPES, 6 KCl, 2 NaCl, 0.5 EGTA, 4MgATP, 0.3 NaGTP,105 Na phosphocreatine. Series resistance (*R*s), typically 17-25 MΩ prior to compensation, was compensated 60%–80%. Bridge balance was used in current clamp. Input resistance (*R*in) and *R*s were monitored in each sweep. Cells were discarded if membrane potential (Vm) at break-in was >-60 mV, *R*in was < 75 MΩ, residual uncompensated Rs was >20MΩ, or if *R*s or *R*in changed by >20% during recording. Vm was corrected for a 12 mV liquid junction potential.

L4 was stimulated in the center of a barrel using a bipolar electrode (0.2 ms, constant-current pulses). L2/3 PYR cells were recorded radially above the stimulus site. Eθ was defined as the minimal intensity that evoked an EPSC. Input-output curves were collected with 10s isi, and 5-6 repetitions of each stimulus intensity. EPSCs and IPSCs were quantified by area or peak, 3-23 ms post-stimulus. L4-evoked PSPs were measured from a pre-stimulus baseline Vm of −50 mV, using the “slow clamp” feature of the Multi-clamp (5 s tau).

mEPSCs and mIPSCs were recorded in TTX citrate (1 μM) and APV (100 μM), without QX-314 in the internal, holding at −72 and 0 mV respectively. In each cell, > 200 mEPSCs and > 200 mIPSCs were detected (criteria: > 5 pA amplitude, 10-90% rise time and peak latency < 2.5 ms) and analyzed using TaroTools (Taro Ishikawa, Jikei University School of Medicine, Japan). Cell-attached spiking was measured using loose-seal recordings in voltage clamp, with holding current at 0 pA. Intrinsic excitability was measured in glutamate and GABA-A synaptic blockers (in μM: 100 APV, 10 NBQX, 3 gabazine).

## Parallel conductance model

Synaptically evoked changes in Vm (ΔVm) were predicted from L4-evoked EPSCs and IPSCs at 1.4 x Eθ using a parallel synaptic conductance model, implemented in Matlab. For each cell, we first calculated the baseline-subtracted mean EPSC at −72 mV and mean IPSC at 0 mV. G_ex_ and G_in_ waveforms were calculated as G = I/(V_hold_-E_rev_), with E_ex_ = 0 mV and E_in_= −72 mV. G_ex_ and G_in_ were constrained to be non-negative and were smoothed (Savitzky-Golay, 1-ms window). Next, we predicted ∆Vm from G_ex_ and G_in_ using the parallel conductance equation (Wehr and Zador, 2003):

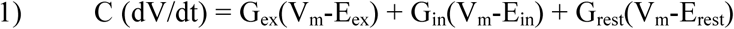

C was 180 pF, which was the average membrane capacitance measured across our genotypes. G_rest_ was defined as 1 / R_input_, where R_input_ was the average input resistance measured for that genotype in current clamp recordings (**Table S4**). We simulated ΔVm for cells at E_rest_ = −50 mV, in order to estimate the effect of feedforward synaptic input on V_m_ as a cell approaches spike threshold. V_m_ was calculated by integrating Eq. 1 from a starting value of V_m_=-50 mV with 0.1 ms time resolution, using Euler’s method^37,57^. Separate analysis was run using Erest = −70 mV.

## In vivo recordings

Adult male mice were anaesthetized with urethane and chlorproxithene (1.3 g/kg and 0.02 mg in saline). Recording was not blind to genotype. A 2 mm craniotomy was made over S1. The mouse was fixed via a head post and the whiskers inserted into piezoelectric actuators. Body temperature was maintained at 36.5° C. Supplemental urethane was provided as needed. The C1, D1, C2 or D2 column was localized by intrinsic signal optical imaging or electrode mapping. A NeuroNexus recording probe (16 or 32 channel) was inserted radially into the target column via a small durotomy. The probe was advanced into L2/3, allowed to settle until stable activity was observed for 30 min, and L2/3 units were recorded. Then the probe was advanced to L4 and allowed to settle again before recording.

Recording location was confirmed either by (i) DiI labeling that was recovered in cytochrome oxidase stained sections that show whisker column boundaries, or (ii) by multi-unit tuning recorded in L4. L2/3 and L4 were defined by microdrive coordinates as 100 - 413 μm and 413-588 μm below the pia^58^. 1 recording site per layer was typically recorded.

### Whisker Stimuli

Calibrated piezo deflections were applied to the column-associated whisker (CW) and the 8 adjacent surround whiskers (SWs) in a 3 × 3 grid, using custom software in Igor Pro. Each whisker deflection was a ramp-hold-return (4 ms – 100 ms – 4 ms). 1.7° deflections were typically used for receptive field mapping. To measure velocity response curves, the CW was deflected at 0.6, 1.1, 1.7, 2.3, 2.9, 3.4, and 4°, with amplitude and velocity co-varying. 75-100 repetitions of each stimulus were presented at each recording site (2-2.5 s isi).

### Analysis

Recordings were amplified and bandpass filtered (Plexon Instruments PBX2/16sp-G50, × 1,000 amplification, 0.3-8 kHz bandpass) and digitized at 31.25 kHz. Noise was reduced by common average referencing. Negative-going spikes were detected using an amplitude threshold (2.8-5 s.d. of background activity), followed by a shadow period of 0.66 ms after each threshold-crossing. 1.5-ms waveforms were clipped for spike sorting. Spike clustering used UltraMegaSort2000 (Hill et al., 2011) and was blind to genotype. Clusters were excluded if they had < 600 spikes, >0.8% refractory period violations (inter-spike interval < 1.5 ms), or > 30% missed spikes (based on Gaussian fit of detected spike amplitudes relative to the detection threshold). FS and RS units were separated using a spike duration criterion of 0.55 ms peak-to-trough time.

### Optogenetic identification of PV interneurons

To validate the spike duration criterion for FS units, we performed a separate series of experiments in which we optogenetically tagged PV interneurons in vivo and identified their spike waveform characteristics. These were performed in adult PV-Cre::ChR2 mice. Recording methods were exactly as described for the main *in vivo* experiments. Once the recording electrode was inserted into S1, we delivered 1-2 ms blue laser flashes (443 nm, 40 mW, CrystaLaser DL445) via an optical fiber (200 μm tip diameter) positioned in air 5 mm above the pial surface. Laser output power was adjusted for each recording site to achieve robust short-latency spiking responses from a subset of units. Units were identified as PV neurons if they exhibited light-evoked spiking at 2-5 ms latency after laser onset with a firing rate 10 standard deviations greater than their baseline firing rate.

#### Quantitative and Statistical Analysis

For slice physiology data, statistical analyses were performed in Prism 7.0 (GraphPad). At least 2 mice and 2 separate litters were used for each measurement. Non-Gaussian data were either log-transformed for parametric testing, or nonparametric tests were applied, as specified in Results. 2-tailed tests were used, with *α*= 0.05. Values in the text are mean ± SEM. Experiments were typically performed blind to genotype and conditions, except in a few cases where more animals of a specific genotype were required to balance the data set. All data analysis was done blind to experimental conditions.

For conductance modeling, predicted PSP peak was quantified as maximum depolarization within 50 ms post-stimulus. Statistical tests are indicated in the figures, and used *α*=0.05. Hypothesized reductions in predicted EPSP or IPSP magnitude (strongly predicted by the voltage-clamp findings in Fig. 1) were tested by 1-tailed KS test. Changes in total predicted PSP were tested by 2-tailed KS test, because no clear prior prediction was available.

For in vivo recordings, analysis was done in Matlab. Spontaneous firing rate was measured in each trial across multiple epochs beginning 0.7 s stimulus offset, which is after whisker-evoked spiking or suppression has subsided. Whisker-evoked spiking was quantified 4-50 ms post-stimulus onset. To determine whether a whisker evoked a significant response from a unit, we computed the probability that a Poisson process with that unit’s mean spontaneous firing would generate the number of spikes measured after whisker deflection, using a binless method. For this test, we used *α*=0.0056 for each whisker (*α*=0.05 / 9 whiskers). Units with significant response to at least 1 whisker were considered whisker-responsive. Whisker receptive field size was the total number of whiskers to which a unit was significantly responsive. The ‘best whisker’ (BW) was defined as the whisker evoking numerically the greatest number of spikes.

Tuning width, tuning accuracy and response latency were calculated only for whisker-responsive units. Response magnitude (e.g., in the velocity response curve) was computed across all single units, including those that were not significantly responsive. Latency was calculated from all combined spikes evoked by significant whiskers, as the earliest time bin at which evoked firing rate exceeded spontaneous firing rate modeled as a Poisson process (*α*=0.05). Jitter was calculated as the standard deviation of spike times 4-50 ms post-stimulus, measured across all whiskers within a unit’s whisker receptive field. Tuning sharpness was defined as the firing rate evoked by the BW divided by the sum of the BW-evoked firing rate plus mean firing rate to all immediately adjacent whiskers. Response latency, jitter and unit depth were normally distributed and genotype differences were evaluated by 2-tailed t-test (*α*=0.05). Velocity response curve data from all units of a given genotype were combined and fit to a sigmoid function using nonlinear regression using the ‘fitnlm’ MatLab function, using the bisquare robust weight option. For VRC fits, statistical differences between genotypes in parameter values were determined by t-test with *α*= 0.007, reflecting Bonferroni correction of total *α*=0.05 across 7 different deflection velocities within the VRC. All other statistical comparisons were made by permutation test with *α*= 0.05. Spike synchrony was calculated from cross-correlograms generated with Matlab’s xcorr() function, using 0.5 ms bin size and ‘coeff’ normalization to remove effects of firing rate. Synchrony was calculated as the mean cross-correlation value over ±10 ms, excluding 0 and ±0.5 ms bins where the shadow period during spike detection prevented simultaneous spikes from being recorded.

